# Predicting Tumor Mutational Burden from UHF-Dielectrophoresis Crossover Frequency

**DOI:** 10.1101/2024.11.05.622085

**Authors:** Héloïse Daverat, Nina Blasco, Sandrine Robert, Arnaud Pothier, Amandine Rovini, Marie Boutaud, Charlotte Jemfer, Claire Dalmay, Fabrice Lalloué, Karine Durand, Thomas Naves

## Abstract

Tumor Mutational Burden (TMB) has emerged as a crucial biomarker to guide patient eligibility for immunotherapy. However, whole exome sequencing, the gold-standard method for TMB measurement, remains limited in accessibility due to its high costs, operational complexity, and lengthy processing times. To address these limitations, we investigated whether Ultra-High-Frequency (UHF) technology could serve as a novel approach to assess TMB by analyzing the crossover frequencies or electromagnetic signature (EMS) of cancer cells on a lab-on-a-chip biosensor, integrating microfluidics and dielectrophoresis.

In a panel of 12 cancer cell lines with varying TMB levels, we observed that EMS showed an upward shift correlating with higher TMB, particularly in solid tumor cell lines. This finding suggests a potential relationship between TMB and EMS. To further explore this hypothesis, we artificially increased mutation levels by treating cells with the highly mutagenic compound N-ethyl-N-nitrosourea (ENU). Results showed that EMS captured significant TMB variations in ENU-treated cells with enhanced proliferative capacity compared to their parental counterparts. These results underscore the importance of matched control samples for reliable EMS measurements. Altogether, our findings highlight the potential of EMS to detect TMB variations associated with proliferative activity, a key hallmark of cancer cells, thereby enabling a more precise stratification of cancer cells.

**Highlights:**

- We propose a new biosensor to improve patient stratification for ICI eligibility
- High frequency fields and dielectric spectroscopy can estimate TMB in cancer cells
- Significant changes in UHF-DEP signatures correlate with varying TMB levels
- UHF-DEP provides a cheap, rapid and label-free method to predict ICI response
- Offers an innovative and complementary marker to conventional diagnostic approaches

**Graphical abstract:** 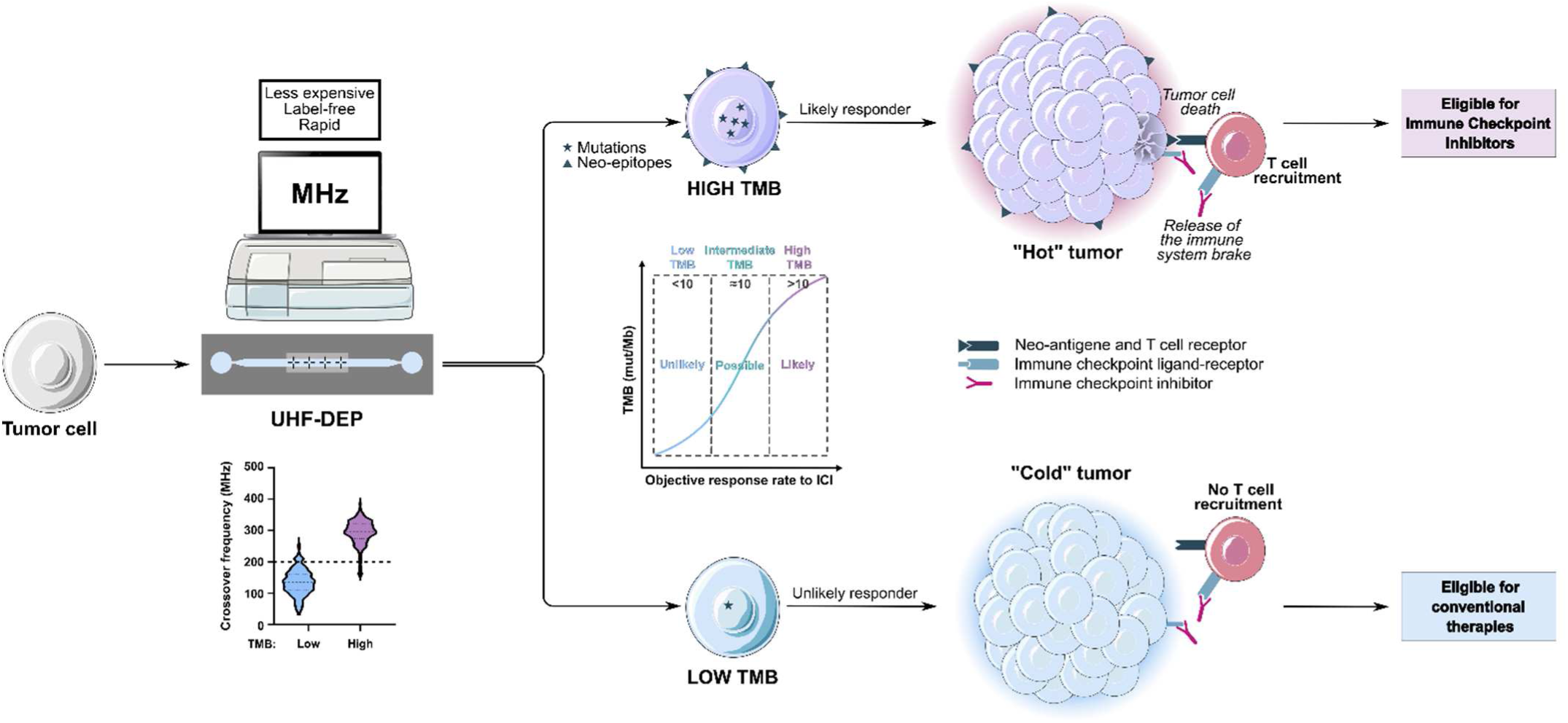

## INTRODUCTION

In the past two decades, immune checkpoint inhibitors (ICI) like anti-PD-1 (Programmed cell Death protein 1) and anti-PD-L1 (Programmed Death-Ligand 1) agents, have transformed the therapeutic landscape of many solid and hematologic malignancies by providing durable responses and improved tolerance (Tan et al., 2020; Shiravand et al., 2022; Salik et al., 2020). These therapies have become central to cancer treatment, especially for patients whose tumors exhibit high PD-L1 expression (Butterfield and Najjar, 2024; Chen et al., 2020). However, although PD-L1 is a relevant biomarker for predicting response to ICI, patients with low PD-L1 expression may also respond favorably to these therapies (Garon et al., 2015; Socinski et al., 2018; Morihiro et al., 2019; Mok et al., 2019). Notably, the FDA-approved (Food and Drug Administration) monoclonal antibody pembrolizumab (Keytruda^®^), which targets PD-1, has demonstrated efficacy in patients with low PD-L1 expression (Wakelee et al., 2023). Conversely, up to 15-40% of patients may fail to respond to immunotherapy despite high PD-L1 expression (Berghmans et al., 2020). If many mechanisms may be responsible for this immunotolerance (Chen and Mellman, 2017), all are far from being elucidated. This variability in response highlights the urgent need for more accurate, robust, and comprehensive biomarkers to better guide ICI-based treatment decisions (Makuku et al., 2021).

Tumor mutational burden (TMB) has attracted a great deal of interest in recent years, with an exponential number of data and publications in the literature. In particular, TMB has emerged as a crucial biomarker for predicting response to immunotherapy (Chalmers et al., 2017; Choucair et al., 2020; Jardim et al., 2021; Aggarwal et al., 2023). TMB measures the total number of non-synonymous somatic mutations per megabase within the coding regions of a tumor genome. Tumors with high TMB, defined as ≥10 mutations/megabase (Mut/Mb), are more likely to respond to immunotherapy (Hellmann et al., 2018; Marabelle et al., 2020a; Litchfield et al., 2021; Gutierrez et al., 2023). Indeed, because mutations generate neoantigens or tumor-specific antigens, which the immune system recognizes as foreign, an anti-tumor response is triggered when ICI release the natural brakes on the immune system (Jardim et al., 2021; Xie et al., 2023). An accurate assessment of TMB can therefore aid in selecting patients who are most likely to benefit from immune checkpoint inhibitors (ICI). However, despite its potential, TMB measurement with the current gold standard, whole exome sequencing (WES), is often costly, resource-intensive, and time-consuming for routine clinical application (Sha et al., 2020).

To address these limitations, we explored the potential of Ultra-High-Frequency (UHF) technology could serve as a novel approach to assess TMB by analyzing the crossover frequencies of cancer cells (Pethig et al., 2010). This approach utilizes on a lab-on-a-chip biosensor that integrates microfluidics and dielectrophoresis (DEP) (Hjeij et al., 2016). The UHF-DEP measures the electromagnetic signature (EMS) of cells by applying a high-frequency, non-uniform field that induces movement in cells suspended in solution. While UHF-DEP does not directly measure mutational burden, it is tempting to speculate that UHF-DEP can be used to analyze various properties indirectly linked to the cells mutational status. Indeed, by altering the DNA sequence, mutations can lead to changes in genome and chromatin structures that disrupt gene expression or to changes in the genes, RNAs and proteins they encode. In particular, non-synonymous mutations cause either a loss or gain of protein function, triggering profound changes in cellular phenotype and behavior. These mutations can lead to alterations in cell signaling pathways, metabolic processes, and even cell cycle regulation and proliferation, key hallmarks of cancer cells (Hanahan and Weinberg, 2011), all of which may contribute to increased tumor aggressiveness therapy resistance.

Here, we propose a model in which cell EMS measured by UHF-DEP could indirectly provide insights into the genomic features of cancer cells. This study potentially opens new avenues for exploring the relationships between genotype, phenotype and biophysical properties, suggesting that EMS indices could be used to determine patient eligibility for anti-cancer treatments. Our findings underscore the potential of EMS to identify high-TMB in solid tumor cell lines with increasing TMB and aggressiveness, thereby supporting more accurate stratification in cancer treatment. Future research will aim to validate these EMS-TMB relationships across a broader range cancer cell lines and tumors, including matched healthy counterparts, to strengthen of EMS’role as a predictive biomarker in oncology.

## MATERIAL AND METHODS

### Tumor samples

Patient DNA for TMB analysis were obtained from the Centre de Biologie et de Recherche en Santé (CRBioLim), as part of project 2023-027, in strict compliance with current regulations governing the use of biological samples for research purposes.

### Cell culture

The four hematological cancer cell lines, Jurkat, MEC-1, OCI-LY10 and TH-P1 were cultured in RPMI 1640 medium (Gibco) supplemented with 10% fetal bovine serum (FBS), 1% antibiotics (penicillin and streptomycin) and 0.1% 2-mercaptoethanol (Gibco). The eight solid cancer cell lines, A549, H1975, U87-MG, DAOY, MEL-5, MEL-28, SW480, and SW620 were cultured under adherent conditions in Dulbecco’s Modified Eagle Medium (DMEM) GlutaMAX (Gibco, Thermo Fisher Scientific, USA), supplemented with 10% FBS (IDbio, France), 1% antibiotics (penicillin and streptomycin, at 100 U/mL and 100 μg/mL respectively) (IDbio). All cell cultures were maintained at 37°C, in a humid atmosphere with 5% CO_2_. Cells were cryopreserved by freezing at −80°C in complete medium containing 10% DMSO (Sigma, France).

### ENU treatment

H1975 and U87-MG cell lines were exposed to 100 μM of the mutagenic chemical agent N-ethyl-N-nitrosourea (ENU, Sigma-Aldrich) for 24 hours to induce an increase in their mutational load. This treatment was repeated 6 times, with a one-week interval between each treatment cycle.

### Dead cells removal

Live cell sorting was performed using the Dead Cell Removal Kit (Miltenyi Biotec, Germany) and MS Columns (Miltenyi Biotec), following the supplier’s recommendations. Cell viability was systematically checked before and after sorting using the LUNA-II™ automatic cell counter (Logos Biosystems, South Korea) and trypan blue staining (Sigma-Aldrich).

### Doubling time calculation

Cells were seeded at a known density and counted using the LUNA-II™ automated cell counter (Logos Biosystems, South Korea). After 48 hours, the cells were detached and counted to determine the final cell density. The doubling time (DT) was calculated using the formula:

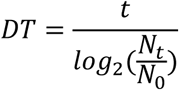

where t is the time interval (48 hours), N_t_ is the final cell count, and N_0_ is the initial cell count.

### Ultra-High Frequency Dielectrophoresis

Biological cells are polarizable entities that, when exposed to a non-uniform electric field, experience forces that enable their manipulation. This force, called the dielectrophoretic force (F_DEP_), depends on the cell’s intracellular dielectric properties, specifically conductivity and permittivity. At higher frequencies, the influence shifts predominantly to the intracellular content and its overall dielectric characteristics. This phenomenon has been demonstrated in previous studies, where ultra-high frequency dielectrophoresis was used to characterize and distinguish cancer stem cells by probing their intracellular content (Lambert et al., 2021; Manczak et al., 2019).

### Genomic DNA extraction and qualification

Genomic DNA was extracted using the Maxwell® CSC Blood DNA or Maxwell® CSC Genomic DNA kits for hematological or solid cancer cell lines respectively, on the Maxwell® CSC Instruments PLC (Promega, USA). The concentration of extracted DNA was measured with the Qubit™ BR dsDNA assay kit (Invitrogen, USA) in a Qubit™ fluorometer (Invitrogen).

### TMB determination by Next-Generation-Sequencing

TMB measurement was performed on 20 ng of genomic DNA with the Oncomine™ Tumor Mutation Load Assay kit (Thermo Fisher Scientific). Quantification of the libraries was carried out using Ion Library TaqMan™ Quantification kit in the QuantStudio 5 real-time quantitative PCR instrument. Libraries were loaded on a Ion 540 chip with the Ion Chef™ Instrument sequencing array preparation system into the Ion S5™ sequencing instrument and sequenced with the Ion S5™ sequencing instrument. Data were analyzed with the online Ion Reporter™ Software.

### Statistical analysis

Statistical analyses, unpaired t-test, Mann-Whitney test, ROC (Receiver Operating Characteristic) curves and AUC (Area Under the Curve), were carried out using GraphPad Prism software (version 10, Dotmatics, USA). Statistical differences were considered significant when the p-value was less than 0.05. Each experiment was performed at least three times independently, ensuring robustness and reproducibility of results.

## RESULTS

### Biosensor to analyze EMS of cancer cells in relation to TMB

We developed a lab-on-a-chip biosensor to measure the electromagnetic signature (EMS) of cancer cells based on their unique intracellular characteristics (Hjeij et al., 2016) (Figure 1A). This device utilizes microfluidics and dielectrophoresis technologies to apply an ultra-high frequency, non-uniform field, which induces movement in cells in suspension (Figure 1A-C). The UHF-DEP signal applied to the cells is generated by a high-frequency source coupled to an amplifier. A splitter is used to provide the same signal to both sides of the sensor. The signal is applied using radio-frequency (RF) probes that contact RF coplanar waveguide (CPW) lines matched to 50 ohms. To prevent standing wave effects and to account for the high impedance of the sensor, 50-ohm (50 Ω) loads are added. The signal applied to the electrodes is monitored on an oscilloscope (Scope) to verify the generated waveform quality. A flow controller that applies pressure to circulate cells in a microfluidic channel in contact with the sensor controls the fluidic part. When a cell is stopped in the middle of the quadrupole a negative DEP (nDEP) signal in high frequency is applied (in the example at 250 MHz in Figure 1D), causing the cell to center itself in the region where the electromagnetic field gradient is weakest (Black square area in Figure 1E). By gradually decreasing the frequency, a slight movement of the cell can be observed, and by decreasing it again until the cell switches to positive DEP (pDEP), the cell will continue to move towards a lateral electrode (white square, Figure 1E). The frequency at which the cellular movement begins is defined as the UHF crossover frequency. For the cell A, the crossover frequency is 230 MHz and 215 MHz for the cell B (Figure 1D). This measurement is repeated on several cells for each sample. While UHF-DEP does not directly measure mutational burden, it can be used to analyze various intracellular properties indirectly linked to the cells mutational status (Figure 1F). Indeed, by altering the DNA sequence, mutations can lead to changes in genome and chromatin structures disrupting spatial genomic organization, transcriptomic and proteomic modifications. In particular, non-synonymous mutations cause either a loss or gain of protein function, triggering profound changes in cellular phenotype and behavior.

**Figure 1:**
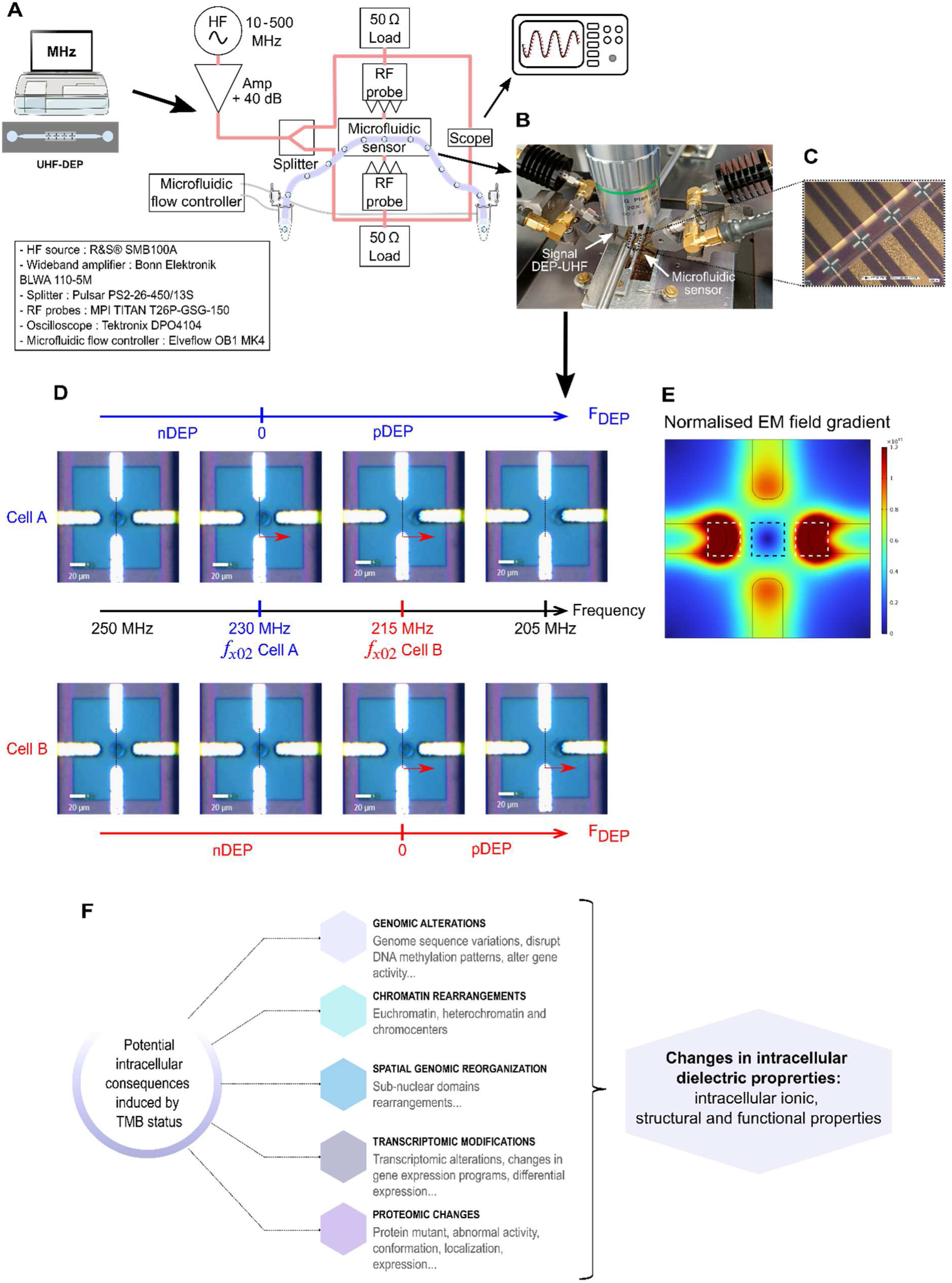
UHF-DEP biosensor for TMB estimation. (**A**) Schematic diagram of the characterization setup where the crossover frequencies measurements are performed: with in red the signal pathway from the high-frequency source (UHF) to an amplifier and to a splitter. The signal is applied using radio frequency (RF) probes. The signal applied to the electrodes is monitored on an oscilloscope (scope) to verify the generated waveform quality. The fluidic part in violet is controlled by a flow controller that applies pressure (in grey) to circulate cells in a microfluidic channel in contact with the sensor. All the references of the setup devices are represented on the left rectangle. (**B**) Photograph of the electromagnetic (EM) sensor under the probes and the microscope. The black circular components represent the 50-ohm loads. (**C**) Zoom on the sensors implemented in the microfluidic channel. This illustrates the coplanar waveguide (CPW) lines that connect multiple quadrupole sensors. (**D**) Measurement of crossovers frequencies with one sensor: dielectrophoretic force (F_DEP_) response of a cell A in blue under an UHF applied signal for frequencies between 250 and 205 MHz. The crossover frequency f_xo2_ is measured at 230 MHz by passing from a negative force (nDEP) to a positive and attractive force (pDEP). And dielectrophoretic force response of a cell B in red under an UHF applied signal for frequencies between 250 and 205 MHz. The crossover frequency f_xo2_ is measured at 215 MHz. (**E**) Numerical simulation of the biased quadrupole in a non-uniform electric field (COMSOL Multiphysics®). The scale color corresponds to the normalised electromagnetic (EM) field gradient intensity 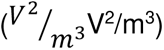. The black square represents the weakest EM field gradient area. The white squares represent the strongest EM field gradient areas, where the cells will be attracted. (**F**) Impact of TMB status on dielectric intracellular properties and their detection through EMS. This schematic illustrates the various intracellular consequences associated with TMB status, including genomic alterations, chromatin rearrangements, spatial genomic reorganization, transcriptomic modifications, and proteomic changes. These alterations can modify intracellular dielectric properties, leading to variations in EMS that can be detected by the UHF-DEP biosensor, highlighting its potential utility in assessing TMB-related cellular characteristics.

Altogether, these consequences of DNA damage suggest investigating whether EMS can indirectly provide insights into the TMB status of cancer cell lines.

### Analysis workflow development: focus on maintaining cell viability

Although EMS depend on several factors, cell viability is a key variable affecting measurements. To avoid these biases, we designed a workflow comprising three steps (step 1 to 3) to eliminate and preserve viable ones before assessing the EMS and TMB measurements of the selected cancer cell line panel (Figure 2A). Indeed, both molecular and cellular changes occurring in dying cells introduce significant biases: (i) alterations in cell structure, and (ii) loss of DNA information following cell membrane disruption. Using a dead cell removal assay, we significantly enriched the proportion of live cells (*p<0.0001*, Figure 2B) prior to suspending them in DEP buffer. This buffer is essential for performing EMS characterization with UHF-DEP (Figure 2C). As expected, EMS values were significantly lower in unsorted cells compared to sorted live cells, likely due to the influence of cell death (*p<0.0001*, Figure 2C). This buffer called DEP, was especially designed to be compatible with both the conductivity (anionic) required for EMS analysis and the physiological osmolarity of cells. Likewise, we analyzed the safety of the DEP buffer concerning cell viability throughout the entire handling process. We observed that cell viability decreases by more than 10% (*p<0.0001*) after 90 minutes of incubation in a time-course study (Figure 2D). To comply with this requirement, the cell handling time after sample preparation (Figure 2A, step 2) was limited to a maximum of 90 minutes to ensure that cell death remained below 10% during EMS characterization (Figure 2A, step 3).

**Figure 2:**
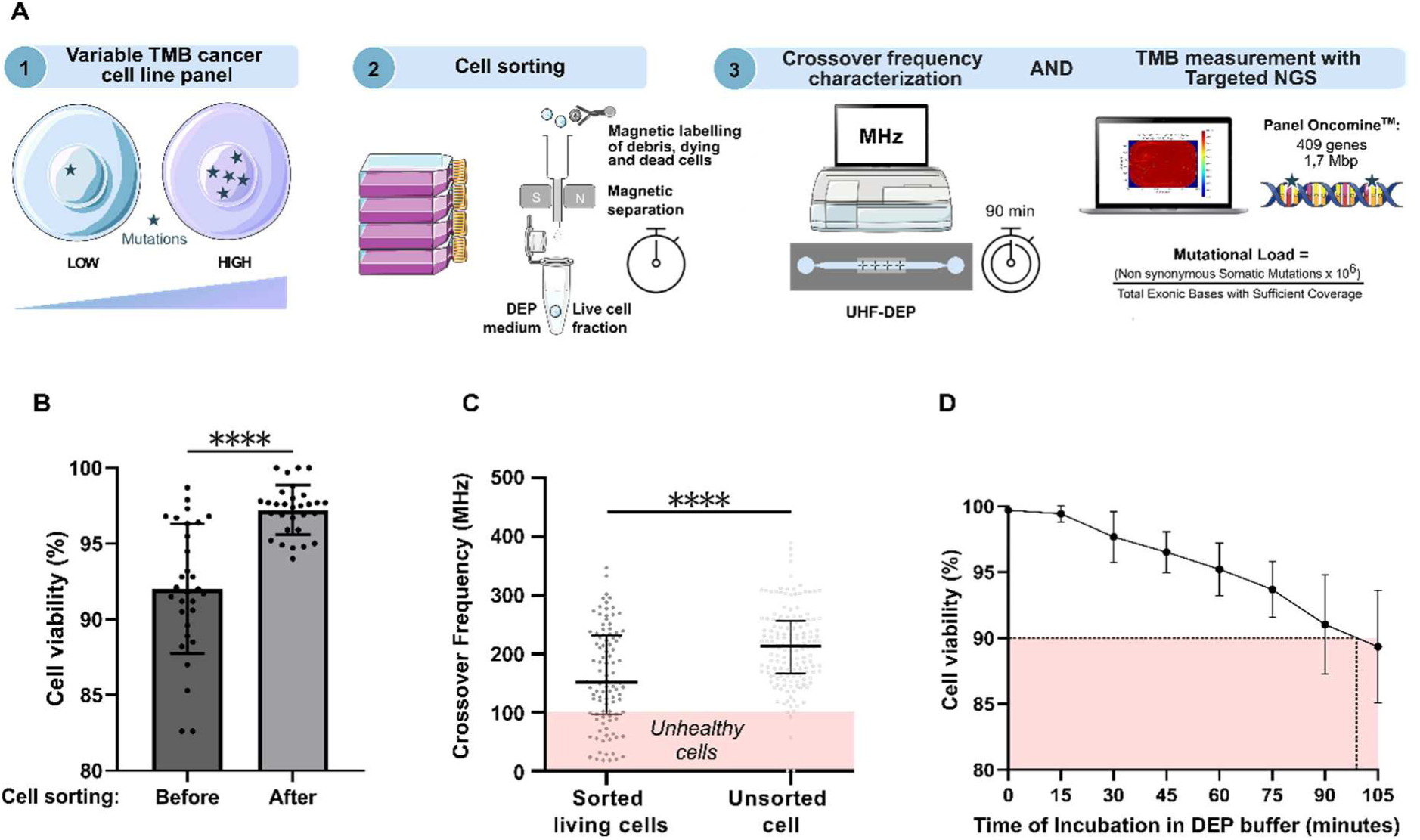
Analysis workflow for ensuring cell viability in reliable EMS and TMB assessment. (**A**) Workflow for assessing EMS and TMB in cancer cell lines. This schematic illustrates the three-step analysis process. Step 1 outlines the selection of a cancer cell line panel characterized by variable TMB levels, ranging from low to high. Step 2 details the sample preparation process, where cells are consistently cultured and prepared using standardized and normalized methods. This includes magnetic labeling and cell sorting, resulting in a live cell fraction suspended in DEP buffer. Step 3 depicts the characterization of crossover frequency using UHF-DEP, conducted over a 90-minute period following the initiation of DEP incubation. TMB measurement is performed using the Oncomine™ targeted NGS panel, with the mutational load calculated by normalizing non-synonymous somatic mutations against the total exonic bases that have sufficient coverage. (**B**) Improved viability of cell population post-sorting. The scatter dot plot with bar illustrates the percentages of viable cells in the population before and after the sorting process across a panel of cancer cell lines (H1975, A549, MEL5, MEL28, SW480, SW620, OCI-LY10, and U87-MG), with a total of 30 samples analyzed. Data are presented as mean values ± standard deviation (SD). Prior to sorting, the mean viability was 92.03% (±4.273), with values ranging from 82.60% to 98.70%. Following the sorting process, the mean cell viability significantly increased to 97.22% (±1.634), with a range of 94.00% to 100%. The enhancement in cell viability following sorting was statistically significant, as determined by an unpaired t-test (**** *p < 0.0001*). (**C**) Crossover frequency of H1975 sorted living cells compared to unsorted cells. The scatter dot plot displays individual crossover frequency measurements in MHz for sorted living cells (N=157) and unsorted cells (N=102). The median crossover frequency for sorted living cells was 213.0 MHz, with an interquartile range (IQR) of 166.0 to 256.0 MHz. For unsorted cells, the median was 150.5 MHz, with an IQR of 96.25 to 231.3 MHz. The enrichment of viable cells significantly increased crossover frequencies, as determined by the Mann-Whitney test (**** *p < 0.0001*). The shaded area indicates the range of crossover frequencies for unhealthy cells. (**D**) Measurement of cell viability in H1975 cells incubated in DEP buffer over time post-sorting. Cell viability remained above 90% for approximately 100 minutes of incubation (n=7). The red shaded area indicates the critical viability threshold set at 90%.

Altogether, this workflow preserves cell viability allowing accurate TMB characterization and improving the reliability of our analyses to better reflect the true biological state of the samples.

### Correlation of tumor mutational burden and crossover frequency in cancer models

Because one major feature of malignant cells lies in the heterogeneity of their mutation levels, we selected a panel of 12 human cancer cell lines representative of both liquid (hematological) or solid cancers (Figure 3A). After removing dead cells, we extracted DNA as templates for TMB measurement before ensuring that each DNA sample met the TMB quality control standards to rule out DNA degradation (data not shown). Then, we analyzed their respective TMB score using the Oncomine™ oncology assay. This panel offers the possibility to use a tumor-only workflow without the need for a matched normal sample, similar to Foundation Medicine (FMI). Indeed, cancer cell lines lack matched normal cell counterparts. Hence, before to assess the accuracy of the Oncomine™ panel we firstly used as templates DNA from two different solid tumor samples (Tumor A and B) for which FMI already assessed TMB (Figure 3B). The Oncomine™ assay revealed TMB level trends like to those observed with FMI, with score differences attributed to variations in gene types, quantity, and sequencing coverage. Following these crucial steps, we assessed TMB in cancer cell lines and obtained values ranging from 3.69 to 114.31 Mut/Mb (Figure 3A). These results indicate broad variability in the mutation burden among the different cell lines used, providing a range of TMB values. This range serves as a valuable reference for comparing TMB across samples, highlighting differences that could influence EMS. Based on this, we first analyzed solid cancer cell lines, which harbor EMS values ranging from 147 to 284.5 MHz (Figure 3C). The box plot shows cell lines in ascending order of TMB, ranging from 3.69 to 17.97 Mut/Mb. A crossover frequency near 210 MHz (grey dashed line) appears to differentiate two populations based on the FDA-defined TMB cutoff of 10 Mut/Mb: high (purple) and low (light blue) (Figure 3C). H1975 and MEL-28 cells have adjacent TMB value to the cutoff of 10 Mut/Mb (respectively 9.56 vs 11.66 Mut/Mb) and EMS values (213 vs 212 MHz), positioning them centrally within the model (Figure 3C). When cell lines were grouped by the FDA’s TMB cutoff (Low in light blue; High in Purple) EMS values showed a significant increase with higher TMB (*p<0.0001*) (Figure 3D) and a positive *Pearson* correlation (*r=0.56*, *p=0.0314*) (Figure 3E). Conversely, hematological cancer cell lines, which harbor the higher TMB of 114.31 Mut/Mb, particularly Jurkat cells (Figure 1A), compared to others, with TMB values ranging from 5.41 to 7.97 Mut/Mb, no significant variation were observed in EMS measurements (Figure 3F). Indeed, difference in crossover frequencies remained around 10 MHz (262 to 272 MHz) (Figure 3G), with no significant correlation between EMS and TMB (Figure 3H).

**Figure 3:**
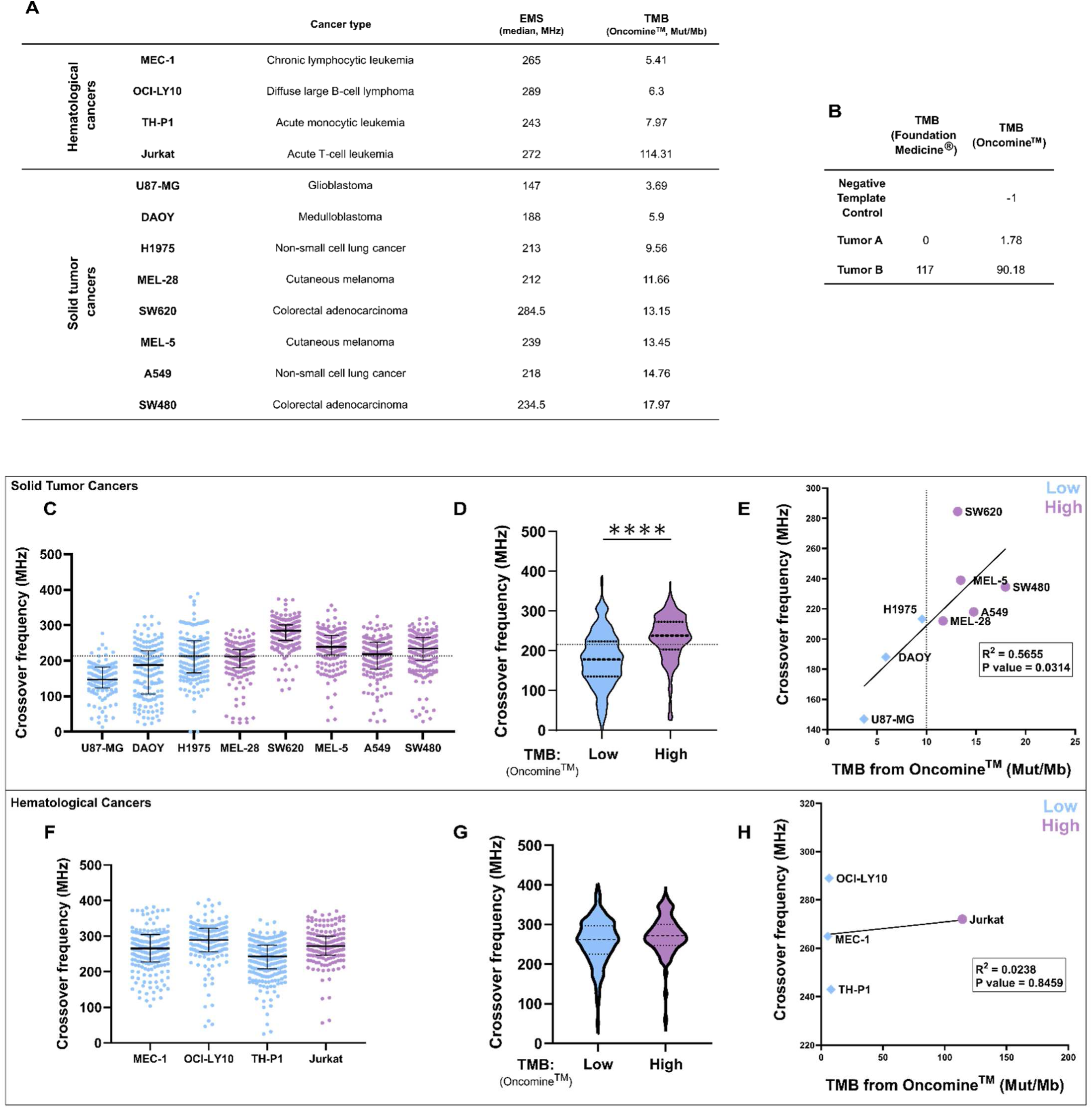
Positive correlation between EMS and TMB in solid cancer cell lines. (**A**) EMS and TMB across selected panel cell lines. This table lists the median EMS values (in MHz) and corresponding TMB scores (using the Oncomine™ assay, in Mut/Mb) for selected hematological and solid tumor cancer cell lines. (**B**) Validation of our TMB calculation method. This table presents TMB values obtained from two assays: Foundation Medicine® (FMI) and Oncomine™. The negative template control serves as an experimental blank, showing a TMB of −1 for Oncomine™. For Tumor A, the TMB is 0 for FMI and 1.78 for Oncomine™, while for Tumor B, the TMB values are 117 for FMI and 90.18 for Oncomine™. The Oncomine™ assay revealed TMB level trends similar to those observed with FMI, this consistency supports the validation of our method for determining TMB. (**C**) The scatter plot displays individual crossover frequency measurements for the solid tumor cancer cell lines included in the panel. Each point represents a distinct measurement, with median values indicated by the horizontal lines and error bars representing IQR. The low TMB group is defined as having TMB values less than 10 Mut/Mb, while the high TMB group includes those with TMB values equal to or greater than 10 Mut/Mb. (**D**) The violin plot compares crossover frequencies categorized by low and high TMB levels based on the Oncomine™ assay for solid tumor cancer cell lines. For the low TMB group, the median crossover frequency was 178.0 MHz (IQR: 135.0 to 223.5 MHz). For the high TMB group (N=868), the median was 238.0 MHz (IQR: 203.0 to 273.0 MHz). The significant difference in crossover frequencies between low (blue) and high (purple) TMB groups was assessed using an unpaired t-test (**** *p < 0.0001*), indicating that higher TMB is associated with increased crossover frequency. (**E**) Linear regression of the relationship between crossover frequency and TMB in solid tumor cancers models. Data points represent SEM median and TBM for each cell line. A vertical dotted line indicates the TMB cutoff of 10 Mut/Mb. The linear regression analysis demonstrates a positive correlation between TMB and crossover frequency (R² = 0.5655; *p value* = 0.0314), indicating statistical significance. This suggests that increased TMB is associated with higher crossover frequencies in the tested solid tumors. (**F**) The scatter plot displays individual crossover frequency measurements for the hematological cancer cell lines. (**G**) The violin plot compares crossover frequencies categorized by low and high TMB levels for hematological cancer cell lines. For the low TMB group (N=612), the median crossover frequency was 262.0 MHz (IQR: 225.3 to 297.0 MHz). For the high TMB group (N=171), the median was 272.0 MHz (IQR: 247.0 to 300.0 MHz). The significant difference in crossover frequencies between low (blue) and high (purple). (**H**) Linear regression of the relationship between crossover frequency and TMB in hematological cancer models. Data points represent the median crossover frequency and TMB for each cell line. The linear regression analysis shows no correlation between TMB and crossover frequency (R² = 0.02375; *p value* = 0.8459), indicating no statistical significance. This suggests that variations in TMB do not significantly influence crossover frequencies in the tested hematological cancers.

Altogether, our findings suggest a correlation between TMB and EMS, especially in solid cancer cell line following the FDA-defined threshold of 10 Mut/Mb. Results in hematological cell line suggest that Oncomine™ panel could not be pertinent.

### Relevance of EMS for determining TMB: validation with CCLE data

However, previous TMB variations between Oncomine™ and FMI in patient samples (Figure 3B) highlight the need for careful validation when applying different panels for TMB measurement across various tumor types, particularly in relation to EMS measurements that can further influence the interpretation of TMB. Consequently, we sorted and curated publicly available TMB data from the Cancer Cell Line Encyclopedia (CCLE) (Ghandi et al., 2019) and the Broad Institute via cBioPortal (https://www.cbioportal.org/) (Cerami et al., 2012). Interestingly, in most cases, TMB values from CCLE were higher than those obtained with the Oncomine™ assay, likely due to differences in gene selection, quantity, and sequencing coverage (Figure 4A). A Spearman correlation analysis indicated a strong and significant correlation between experimental TMB values obtained using Oncomine™ panel and those from CCLE (r=0.94, *p<0.0001*) (Figure 4B). Notably, the CCLE data places the intermediate cell line H1975 over the FDA-defined TMB cutoff of 10 Mut/Mb (11.36 Mut/Mb), underscoring the need for careful validation of TMB values depending on the panel used. Hence, using the CCLE-based TMB values to the previous EMS obtained for each cell lines, we also categorized two populations based on the FDA-defined TMB cutoff of 10 Mut/Mb: high (purple in the box plot) and low (light blue in the box plot) (Figure 4C and 4D). Using EMS data grouped by TMB levels, we found that an EMS around 200 MHz significantly distinguishes cells with high TMB from those with low TMB only in solid cancer cell lines (*p<0.0001*) (Figure 4D). However, adding EMS data from hematological cell lines disrupt this model, significantly reducing sensitivity, with the difference becoming only weakly significant (*p<0.05*) (Figure 4D). The ROC curve generated from EMS results of solid cancer cell lines using the FDA TMB cutoff of 10 for low and high TMB, indicates relevant AUC values (Area Under the Curve) (Figure 4E). The dark line represents an AUC of 0.86 for the Oncomine™ panel, while the light blue line indicates an AUC of 0.93 for CCLE, confirming the strong accuracy of EMS in discriminating solid cancer cells based on their TMB (Figure 4E).

**Figure 4:**
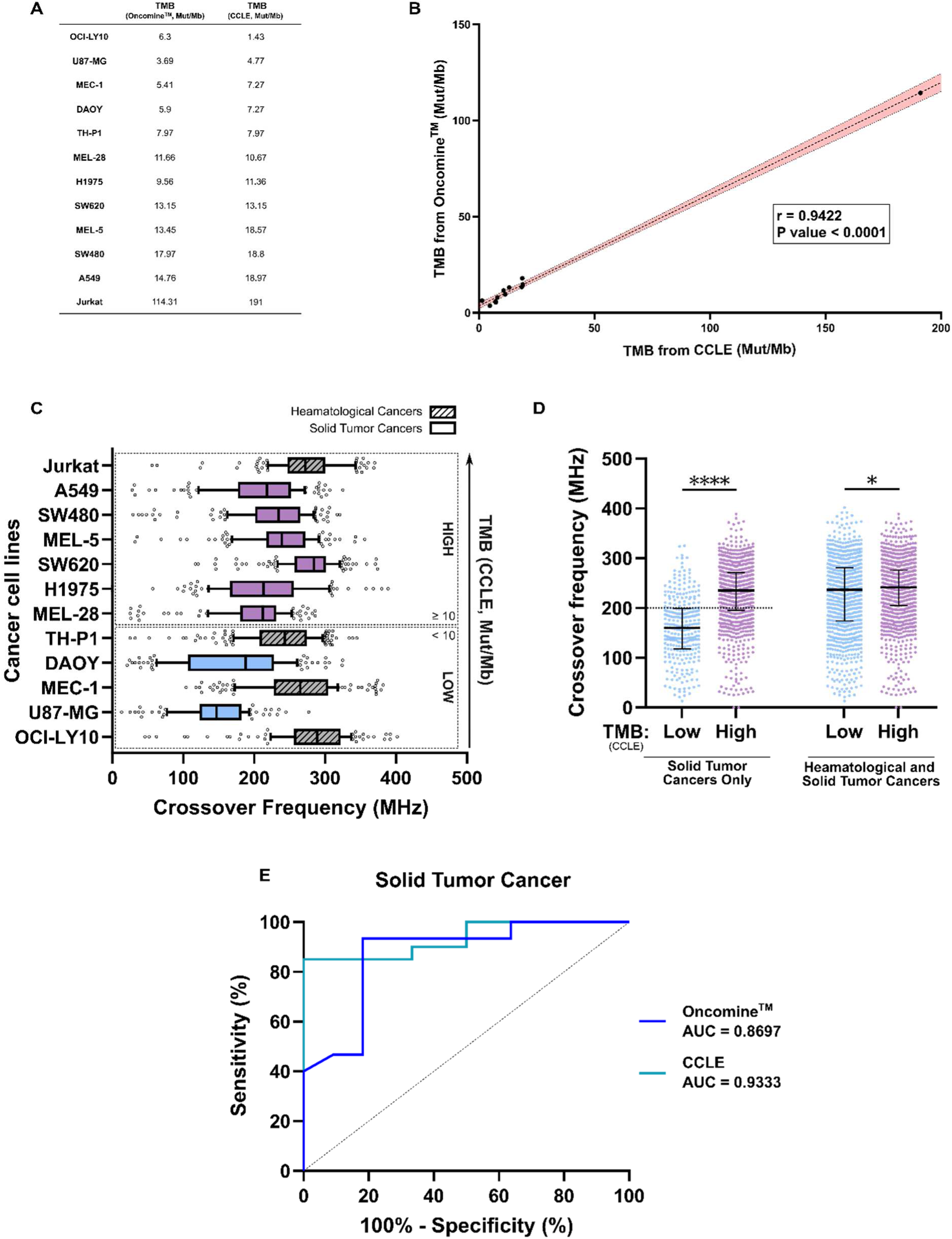
EMS accurately predict TMB in solid cancer cell lines: validation with CCLE data. (**A**) Comparison of Tumor Mutational Burden (TMB) Measurements from Oncomine™ and CCLE. This table presents TMB values (in Mut/Mb) obtained from two assays for various cancer cell lines, comparing results from the Oncomine™ panel with those from the Cancer Cell Line Encyclopedia (CCLE). (**B**) Spearman Correlation of Tumor Mutational Burden (TMB) from Oncomine™ and CCLE. The scatter plot illustrates the correlation between TMB values (in Mut/Mb) obtained from the Oncomine™ assay and those from the Cancer Cell Line Encyclopedia (CCLE). Each data point represents the TMB measurement for a specific cancer cell line. A strong positive Spearman correlation (*r* = 0.9422; *p < 0.0001*, 95% confidence interval: 0.7949 to 0.9846) was observed, indicating a significant relationship between TMB values from both assays. The shaded area represents the 95% confidence interval, with the boundaries indicated by the dotted black lines. (**C**) Crossover frequency in both solid and hematologic cancer cell lines based on TMB from CCLE. The cancer cell lines are arranged along the y-axis, with those above the defined TMB cutoff of 10 Mut/Mb indicated in purple (high TMB) and those below in blue (low TMB). The plot represents crossover frequency as box and whiskers, where the whiskers extend to the 10th and 90th percentiles. (**D**) The scatter dot plot displays crossover frequency measurements categorized by low and high TMB levels, as determined by CCLE data. The left side of the plot shows results for solid tumor cancers only, with the low TMB group (N=300) having a median crossover frequency of 160.0 MHz (IQR: 118.3 to 199.0 MHz) and the high TMB group (N=1025) showing a median of 235.0 MHz (IQR: 195.0 to 271.0 MHz). The right side of the plot includes both hematological and solid tumor cancers, where the low TMB group (N=912) has a median crossover frequency of 237.0 MHz (IQR: 174.0 to 281.0 MHz), while the high TMB group (N=1196) has a median of 241.5 MHz (IQR: 205.0 to 276.0 MHz). The significant difference in crossover frequencies between low and high TMB groups in both categories was assessed using the Mann-Whitney test (* *p < 0.05*; **** *p < 0.0001*), indicating that higher TMB is associated with increased crossover frequency especially in solid tumor cancer cell lines. (**E**) Receiver Operating Characteristic (ROC) Curves for TMB Prediction in Solid Tumor Cancers. The ROC curves illustrate TMB prediction based on data from the Oncomine™ and CCLE assays, with classification applied at a cutoff of 10 Mut/Mb for each panel. The dark blue curve represents the Oncomine™ classification, with an area under the curve (AUC) of 0.8697 (95% confidence interval: 0.7224 to 1.000; *p value* = 0.0015). The cyan curve represents the CCLE classification, with an AUC of 0.9333 (95% confidence interval: 0.8382 to 1.000; *p value* = 0.0016). These results indicate a high level of accuracy for both panels in distinguishing between high and low TMB levels in solid tumor cancers.

Altogether, these findings suggest that EMS can predict indirectly the TMB level in solid cancer cell lines.

### Assessing TMB and EMS in response to evolving mutations

Given previous results suggesting that an EMS cutoff around 200 MHz can predict a TMB threshold above 10 Mut/Mb in solid cancer cell lines, we conducted further analyses to investigate whether EMS could monitor TMB changes in response to accumulating mutations. Due to the absence of matched normal samples for comparison with cancer cell lines, and considering that cancer is an evolving genetic disease, we artificially increased mutation levels in cell lines with either low (U87-MG) or intermediate (H1975) TMB scores. We then monitored changes in EMS, specifically observing whether TMB exceeded the FDA cutoff of 10 Mut/Mb compared to their respective parental cell lines. To test this hypothesis, we treated cells with the highly mutagenic compound N-ethyl-N-nitrosourea (ENU) at a concentration of 100 μM for 24 hours, repeating this cycle multiple times. We arbitrary conducted six consecutive cycles of ENU treatment, tracking EMS as the cells acquired new mutations or affecting their respective TMB (Figure 5A). Likewise, since ENU induces random mutations that may or may not influence cell fate, we assessed whether these hereditary mutations confer positive or negative advantages by selecting a cancer cell population with an increased proliferative doubling time.

**Figure 5:**
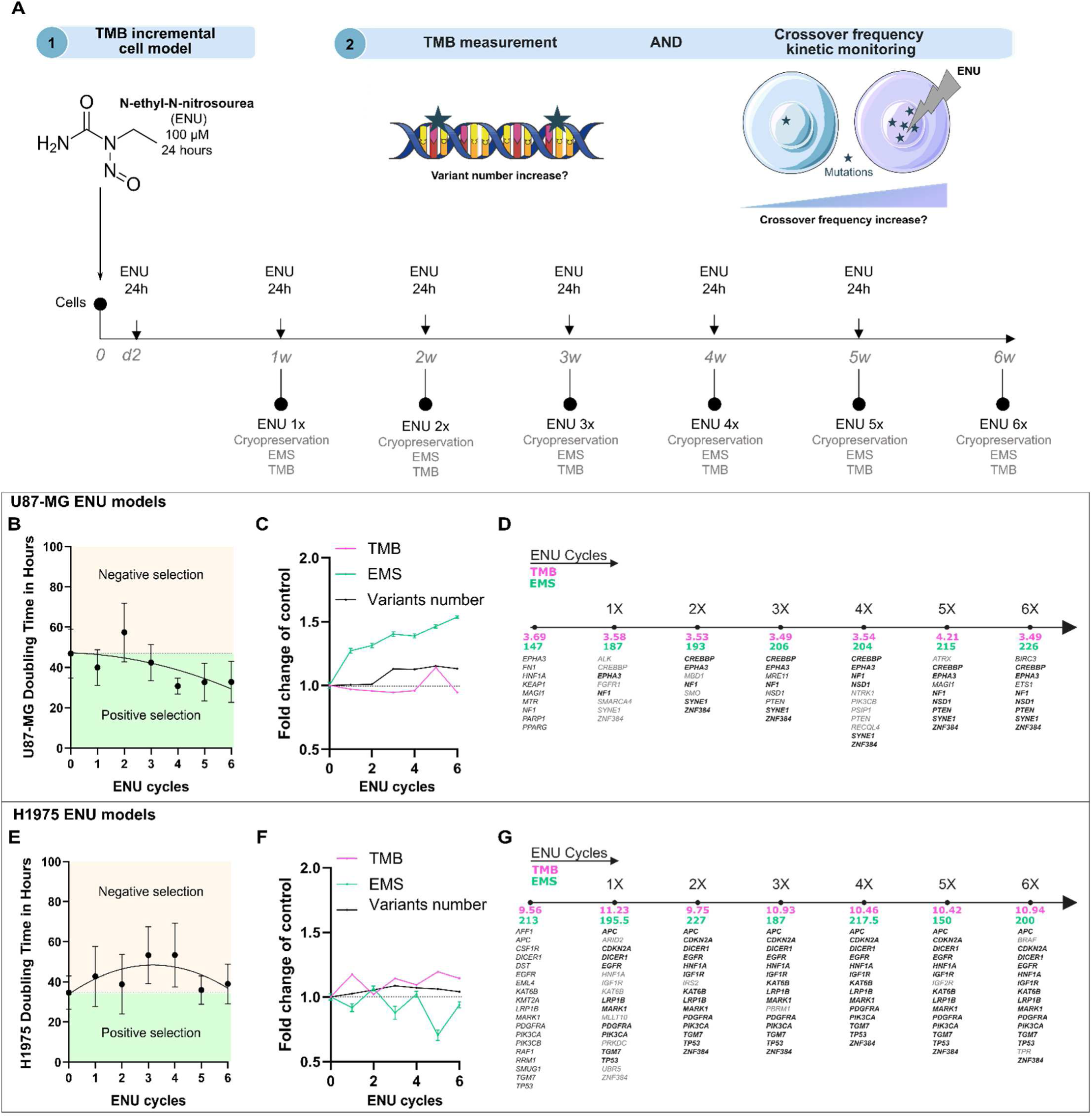
Tracking the Evolution of TMB and Electromagnetic Signatures in Response to Induced Mutations. (**A**) This diagram illustrates the timeline and procedures for inducing mutations in cancer cell lines using 100 μM ENU treatment. Step 1: Mutation Induction. One million cells from the U87-MG and H1975 cell lines are seeded and treated with ENU (100 μM) in complete medium for 24 hours, starting 2 days post-seeding (d2). After ENU exposure, the cells are washed and incubated for 1 week (1w). ENU treatment is repeated for a total of six cycles. Following the last treatment, ENU-treated cells are harvested and divided for further ENU treatments and cryopreservation for subsequent analysis. Step 2: Simultaneous EMS and TMB Analysis. EMS and TMB analyses are performed simultaneously, allowing for the evaluation of the correlation between induced mutations and changes in TMB and EMS. (**B** and **E**) This graph illustrates the doubling time in hours for U87-MG and H1975 cells across six cycles of ENU treatment. The shaded regions differentiate between negative selection (beige) and positive selection (green), illustrating the impact of ENU-induced mutations on the growth dynamics of the cell lines. (**C** and **F**) EMS and TMB data were collected for each cycle of ENU treatment in the U87-MG and H1975 cell lines, along with the number of variants identified. The number of variants corresponds to a specific alteration of the DNA sequence compared to the reference sequence, such as a point mutation, an insertion, a deletion, or another type of genetic change. The analysis of variant and TMB calculation was conducted using Ion Reporter software, patch 5.18, with the “TMB extended’ option”. For each cell line, values were normalized against the control condition (untreated wild-type cells) and expressed as fold change relative to control. (**D** and **G**) Phylogenetic analysis of induced mutations across ENU treatment cycles in U87-MG and H1975 cell lines. Genes that are conserved across the different ENU cycles (1X to 6X) are highlighted in bold, with respective TMB and EMS value annotated. In U87-MG cells, mutations in key genes are involved in tumor suppression and transcriptional regulation, appeared and were conserved across ENU cycles, suggesting a potential selective advantage. In H1975 cells, mutations in oncogenes and tumor suppressor genes were preserved across ENU cycles.

Interestingly, we observed that ENU treatment decreased the doubling time of U87-MG cells, suggesting that newly acquired mutations positively select for a cell population (Figure 5B). In contrast, the doubling time of H1975 cells increased from the early cycles of ENU treatment (Figure 5E). TMB measurements in terms of fold change relative to control in both cell lines reveal opposite effects of ENU, with a rapid TMB increase in H1975 cells (pink curves, Figure 5F) compared to U87-MG cells, which show a general slowdown until reaching a peak at the fifth cycle of ENU (pink curves, Figure 5C). However, the number of variants, indicating DNA sequence variation, increased from the second cycle of ENU in U87-MG cells (black curves, Figure 5C) while remaining with a minimal increase in H1975 cells (black curves, Figure 5F). Interestingly, only U87-MG cells exhibited a continuous increase in EMS throughout the ENU cycles (green curves, Figure 5C). In contrast, the EMS evolution in H1975 (green curves, Figure 5F) cells appeared to depend on the paired ENU cycles, showing a tendency to decrease. Strikingly, U87-MG cells exceeded the EMS cutoff of 200 MHz from the third ENU treatment cycle onward (Figure 5D), although their TMB score remained below the FDA cutoff of 10 Mut/Mb. Likewise, although ENU-treated H1975 cells exceeded 10 Mut/Mb depending on the ENU cycle (Figure 5G), their associated EMS remained below 200 MHz. Analysis of mutation patterns (Figure 5D and 5G) reveals a difference in the heritability of mutations between U87-MG and H1975 cells. In U87-MG, the mutation pattern remains stable from the fifth ENU cycle, while H1975 exhibits an almost complete mutation pattern from the early ENU cycles. Similarly, the difference in the number of mutated genes in H1975 cells may trigger negative selection, as evidenced by their increased doubling time.

Altogether, these results indicate that EMS and TMB are not necessarily correlated and underscore the importance of matched controls. EMS may reflects cell state variation that are influenced by mutation types in TMB, which can impact cell fate through positive or negative selection.

## DISCUSSION

Biomarkers, whether prognostic or predictive of treatment response, form the foundation of personalized medicine, one of the main public health challenges in oncology worldwide. Developing new tools capable of rapidly assessing these biomarkers, to minimize loss of opportunity, and affordably, to reduce costs for society is a key objective of translational and interdisciplinary research to further refine patient stratification. The emergence of immune checkpoint inhibitors (ICIs) has radically transformed treatment in numerous cancers with a growing need to optimize patient selection and develop reliable biomarkers (Rui et al., 2023). TMB is emerging as a potential biomarker and has been FDA-approved to determine patient eligibility for pembrozulimab (Marabelle et al., 2020b). However, significant challenges remain before its widespread and effective adoption, particularly due to the high cost, resource demands, and time requirements of whole exome sequencing, the primary method for measuring TMB, which limits routine clinical application (Sha et al., 2020). Additionally, there are ongoing uncertainties about the applicability of a universal threshold of 10 mut/Mb (Budczies et al., 2024). To address these limitations, we explored Ultra-High-Frequency technology as a novel method to assess TMB by analyzing cancer cell crossover frequencies. Mutations alter DNA sequences, potentially affecting gene expression and protein function, and leading to changes in signaling pathways, metabolism, cell cycle, and proliferation, hallmarks of cancer (Hanahan, 2022). Although UHF-DEP does not directly measure mutational burden, it may enable indirect analysis of mutation-linked properties.

The workflow we developed using UHF-DEP demonstrated its ability to distinguish cells with high TMB, based on a cutoff of 200 MHz. High and Low TMB classification was determined by Oncomine™ and defined as ≥ 10 Mut/Mb according to FDA approval, without requiring the companion diagnostic test performed by FMI. While this approach is applicable to solid cancer cell lines, hematological cells do not follow this rule. Interestingly, we observed a correlation between EMS and TMB across the various solid cancer cell lines analyzed. However, results obtained after treatment with the highly mutagenic agent ENU lead us to make certain recommendations. ENU induces random mutations in the genome, even though this type of mutagenesis does not follow the pathophysiological processes typically seen in oncogenesis. ENU treatment cycles drive dynamic changes in the genome (TMB and variant counts) and variations in EMS. This suggests that TMB might be better considered as a measure of variation between the original cell and the transformed cell, rather than as an absolute value. Further experiments are needed to test this hypothesis, such as inducing oncogenic transformation through a more pathophysiological process or using primary cancer cells directly. In clinical applications, this would imply comparing the EMS of tumor cells to that of their healthy counterparts, which would provide more relevant references. This study underscores the potential of UHF-DEP technology to refine biomarker assessments, particularly for TMB, enhancing patient stratification for immune checkpoint inhibitor (ICI) therapies. By enabling the detection of TMB variations, UHF-DEP could complement existing diagnostic tools, facilitating more personalized and accurate treatment selection in oncology (Figure 6).

**Figure 6:**
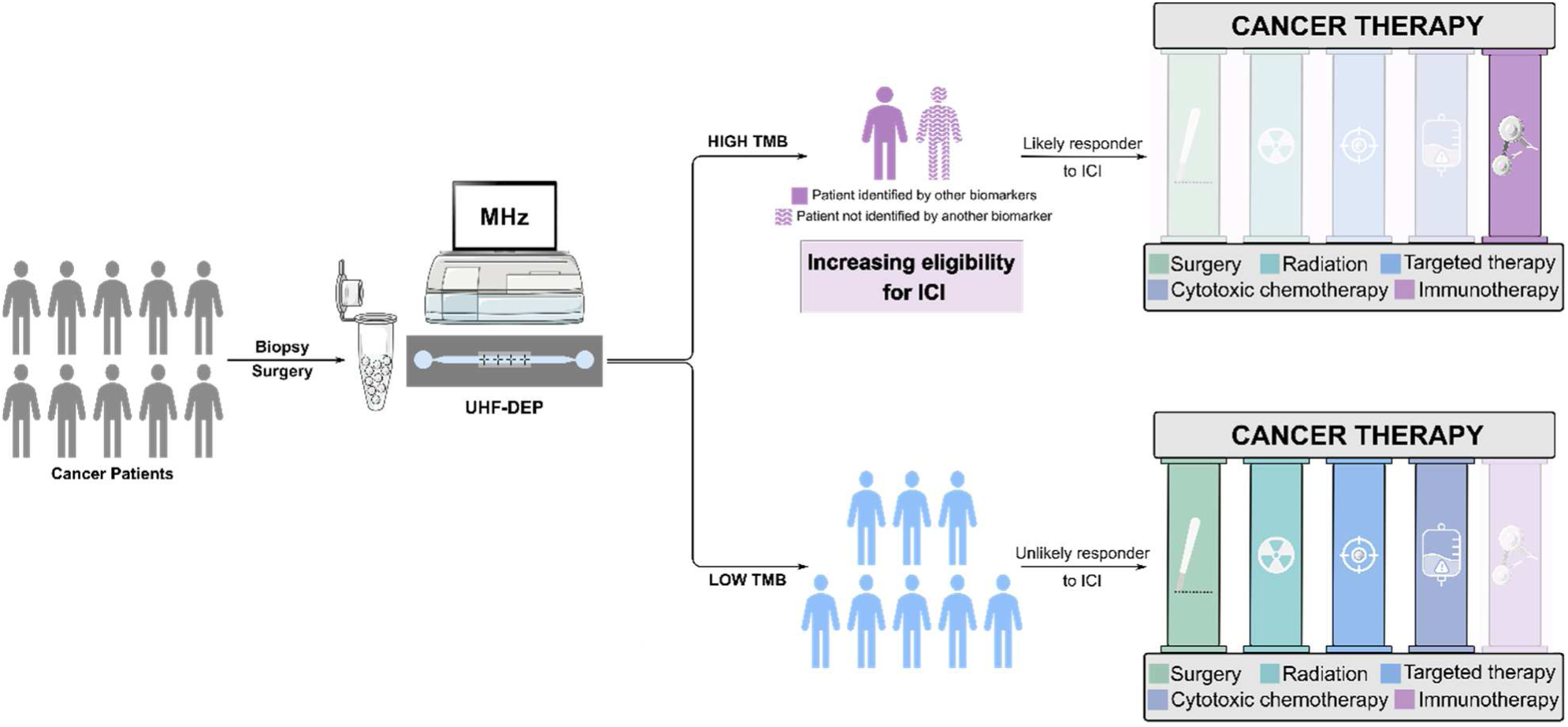
Leveraging UHF-DEP for enhanced therapeutic decision-making in cancer treatment. This schematic illustrates the potential application of the UHF-DEP tool in guiding therapeutic decisions for cancer patients. By characterizing tumor cells obtained from biopsies or surgical procedures, UHF-DEP can effectively discriminate between low and high SEM, which reflect low and high TMB, respectively. The integration of UHF-DEP with existing biomarkers aims to refine patient eligibility for ICI, enhancing the stratification of cancer patients. Those identified with high TMB through UHF-DEP can be prioritized for ICI therapy. In contrast, patients with low TMB may be directed towards alternative cornerstone treatment options, including surgery, radiation, targeted therapies, or cytotoxic chemotherapy. This approach not only optimizes patient outcomes by ensuring that those who are most likely to benefit from ICI are identified and treated accordingly, but it also streamlines therapeutic pathways within the overall treatment algorithm for cancer care, limiting analysis costs per patient and providing rapid responses.

## DECLARATIONS

## Aknowlodgement

This study was supported by the Ligue Contre le Cancer Limousin, Comité d’Orientation sur le Cancer, CHU Dupuytren, Limoges. HD PhD. was supported by the Région Nouvelle Aquitaine. CRBiolim, Centre de Ressources Biologiques, CHU Dupuytren Limoges for patient samples and UF5315, Unité fonctionnelle de support à la recherche translationnelle et innovation en oncologie solide for their research support.

## Author contributions

TN, KD, FL, AP and CD contributed to the conception and design of the study and the study protocol. HD and NB performed the experimental work and performed data analysis. AR proof-read. SR, MB, CJ and AR performed research. All authors read and approved the final version of the manuscript.

## Data availability statement

Data will be made available upon reasonable request to the corresponding author.

